# Antiviral and Virucidal activities of sulphated polysaccharides against Japanese Encephalitis Virus

**DOI:** 10.1101/2020.02.11.944116

**Authors:** Hussin A. Rothan, Rohana Yusof

## Abstract

**Background:** Japanese encephalitis virus (JEV), a member of the family Flaviviridae, causes severe neurological disorders in humans. JEV infections represent one of the most widely spread mosquito-borne diseases, and therefore, it has been considered as an endemic disease. An effective antiviral drug is still unavailable to treat JEV, and current drugs only provide supportive treatment to alleviate the symptoms and stabilize patients’ conditions. This study was designed to evaluate the antiviral activity of the sulphated polysaccharides “Carrageenan,” a linear sulphated polysaccharide that is extracted from red edible seaweeds against JEV replication in vitro.

**Methods and Results:** Viral inactivation, attachment, and post-infection assays were used to determine the mode of inhibition of Carrageenan. Virus titters after each application were evaluated by plaque formation assay. MTT assay was used to determine the 50% cytotoxic concentration (CC50), and ELISA-like cell-based assay and immunostaining and immunostaining techniques were used to evaluate the 50% effective concentration (EC50). This study showed that Carrageenan inhibited JEV at an EC50 of 15 μg/mL in a dose-dependent manner with CC50 more than 200 μg/mL in healthy human liver cells (WRL68). The mode of inhibition assay showed that the antiviral effects of Carrageenan are mainly due to their ability to inhibit the early stages of virus infection such as the viral attachment and the cellular entry stages.

**Conclusion:** Our investigation showed that Carrageenan could be considered as a potent antiviral agent to JEV infection. Further experimental and clinical studies are needed to investigate the potential applications of Carrageenan for clinical intervention against JEV infection.

## Introduction

Flaviviruses infection has been disseminated across many countries in tropical countries [1]. Japanese Encephalitis virus (JEV) infections represent one of the most widely spread mosquito-borne diseases, and therefore, it has been considered as an endemic disease, predominantly occurs in South-East Asia and the Western Pacific regions [2, 3]. The infection is mainly caused by the transmission of JEV to humans through mosquitoes from the Culex genus, especially *Culex tritaeniorhynchus*. The symptom of JEV infection ranges from mild to severe manifestation. Most infected individuals only experience mild symptoms such as fever and headache. However, some (□ 1%) might develop severe clinical symptoms such as high fever, spastic paralysis, and coma [1, 4]. There are approximately 44,000 new cases of JEV with clinical symptoms per year with a 30% mortality rate, while 30-50 % of survivors will suffer permanent neurological impairment [4, 5]. Although vaccines for JEV are available, long term protective effects is still not known, and treatment usually requires at least one booster dose after 12 months of vaccination. This requirement is highly inconvenient and usually leads to low public compliance [6]. An effective antiviral drug is still unavailable to treat JEV, and current drugs only provide supportive treatment to alleviate the symptoms and stabilize patients’ conditions [3]. As such, the development of active antiviral agents for JEV is necessary to address the unmet medical needs.

Natural compounds show considerable antiviral activities through direct binding with viral proteins or cell membrane receptors [7, 8]. Carrageenan is sulphated high-molecular-weight polysaccharides extracted from red edible seaweeds and widely used in the food industry. Chemically, Carrageenan is made up of repeating galactose units joined by alternating α-1,3 and β-1,4 glycosidic linkages. Sulphated polysaccharide was found to be very potent and selective inhibitors of Human Immunodeficiency Virus-1 (HIV-1) properties [9, 10]. These effects are believed to be the result of direct binding to HIV glycoprotein 120 (gp120) on the surface of the viral particle that, in turn, inhibit the entry of HIV-1 into human cells [9]. In this study, we report the antiviral properties and cytotoxicity of Carrageenan towards JEV on WRL68 cells. The mechanisms of the inhibitory effects of Carrageenan were also investigated in order to get a better understanding of Carrageenan as a potential antiviral agent for combating JEV infections.

## Materials and Methods

### Virus and cells

The stocks of JEV were prepared in C6/36 cells, and viral titers were determined *via* plaque assay using Vero cells (ATCC; Rockville, MD, USA). The cells were cultured in complete Dulbecco’s modified Eagle’s medium (DMEM) supplemented with 10% fetal bovine serum (FBS) as a growth medium or 2% FBS as maintenance medium. Iota Carrageenan and heparin were purchased from Sigma (Sigma Aldrich) and dissolved in 5% DMSO. The final concentration of DMSO was lower than 1% in all cell culture-based assays.

### Cytotoxicity assay

Healthy liver cells (WRL68, 1 × 104 cells per well of a 96-well plate) were treated with increasing concentrations (12.5, 25, 50, 100, and 200 µg/ml) of Carrageenan or heparin as a positive control for 72 h. The cytotoxic effect of the test compound was determined based on MTT (3-(4,5-dimethylthiazol 2-yl)-2,5-diphenyltetrazolium bromide) assay, as previously described [11].

### Virus quantification by plaque formation assay

A 10-fold serial dilution of culture supernatant of JEV infected cells was added to fresh Vero cells grown in 6-well plates (1.2× 106 cells) and incubated for 1 h at 37°C. The cells were overlaid with DMEM containing 1.1% methylcellulose. Viral plaques were stained with crystal violet dye after a 5-day incubation period. Virus titers were calculated according to the following formula: Titer (p.f.u./ml) = number of plaques × volume of the diluted virus added to the well × dilution factor of the virus used to infect the well in which the plaques were enumerated [12].

### Evaluation of antiviral activities of Carragennan on WRL68 cell line

The cells were seeded into 24-well plates (1 × 105 cells per well) for 24 h at 37°C and 5% CO2. JEV culture supernatant was separately added to the cells at an M.O.I. of 1, followed by incubation for two h with gentle shaking every 15 min to achieve optimal virus-to-cell contact. The cells were washed twice with phosphate-buffered saline (PBS) after removing the virus culture supernatant. Subsequently, the new complete DMEM containing 25 µg/ml of Carrageenan or heparin was added to the cultures and further incubated for 72 h. Viral titers were evaluated using plaque formation assays.

### Viral inactivation assay

Virus inactivation assay was performed to assess the ability of Carrageenan to inactivate JEV, and prevent subsequent infections directly. WRL68 cells were seeded in 24-well plates (1 × 105 cells/well) for 24 h. The test compound (25 µg/ml) was mixed with the virus culture supernatant (M.O.I of 2) and incubated at 37°C for 1 h and then inoculated onto the WRL68 cells with gentle shaking every 15 min for 2 h. The virus and test compound mixture were then removed, and the cells were washed three times with PBS to remove any residual virus. Then, fresh complete DMEM was added, and the cultures were further incubated for 72 h at 37°C supplemented with 5% CO2. The viral titers of DENV2 and JEV suspension were then determined via plaque formation assays.

### Viral attachment assay

This assay was performed to show the ability of Carrageenan to inhibit DENV2 and JEV entry into host cells. WRL68 cells were grown in 24-well plates (1 × 105 cells/well) for 24 h. Cell culture media were removed and the cells were then washed three times with PBS. New media containing virus and the test compound (25 µg/ml) mixture were separately added and the cells were incubated for one h at 4°C. Subsequently, the media was removed and the cells were washed extensively with cold PBS to remove the unabsorbed virus. The cells were overlaid with DMEM containing 1.1% methylcellulose and incubated for five days. Viral titers were quantified by counting infectious centers.

### Post-infection treatment

The cells were seeded in 24-well plates (1 × 105 cells/well) for 24 h. Cell culture media were removed and the cells were then washed three times with PBS. New media containing virus (M.O.I of 2) was added and incubated at 37°C with gentle shaking every 15 min for two h. The virus was then removed by washing the cells twice with PBS followed by the addition of new complete DMEM containing (25 µg/ml) of the test compound. The cells were then incubated for 72 h. The viral titers of JEV were then determined by plaque formation assays.

### Immunofluorescence assay

Vero cells grown on coverslips were infected with JEV (M.O.I. of 2) in the presence or absence of Carrageenan (10, 20 and 30 μg/ml). At 72 h post-infection, cell monolayers were washed with cold PBS and fixed in methanol for 15 min at -20 °C for cytoplasmic immunofluorescence. Indirect staining was carried out by using anti-JEV mouse antibodies and fluorescein (FITC)-labeled goat anti-mouse IgG [13].

### Statistical analysis

All assays were performed in triplicates, and the statistical analyses were performed using GraphPad Prism version 5.01 (GraphPad Software, San Diego, CA). P values <0.05 were considered significant. The error bars are expressed as ± SD.

## Results

### Determination of potential cytotoxicity of Carrageenan

The potential cytotoxicity effect of Carrageenan was evaluated using WRL68 cells and heparin was employed as a positive control. The 50% cytotoxic concentration (CC50) was higher than 200 µg/ml for both Carrageenan and heparin The maximal non-toxic dose (MNTD) value was approximately 25 µg/ml and showed at least 95% cell viability. Following this, all subsequent in vitro cell culture experiments were carried out using doses of less than 25 µg/ml of the potential compounds (Fig. 1A and 1B).

**Figure 1:**
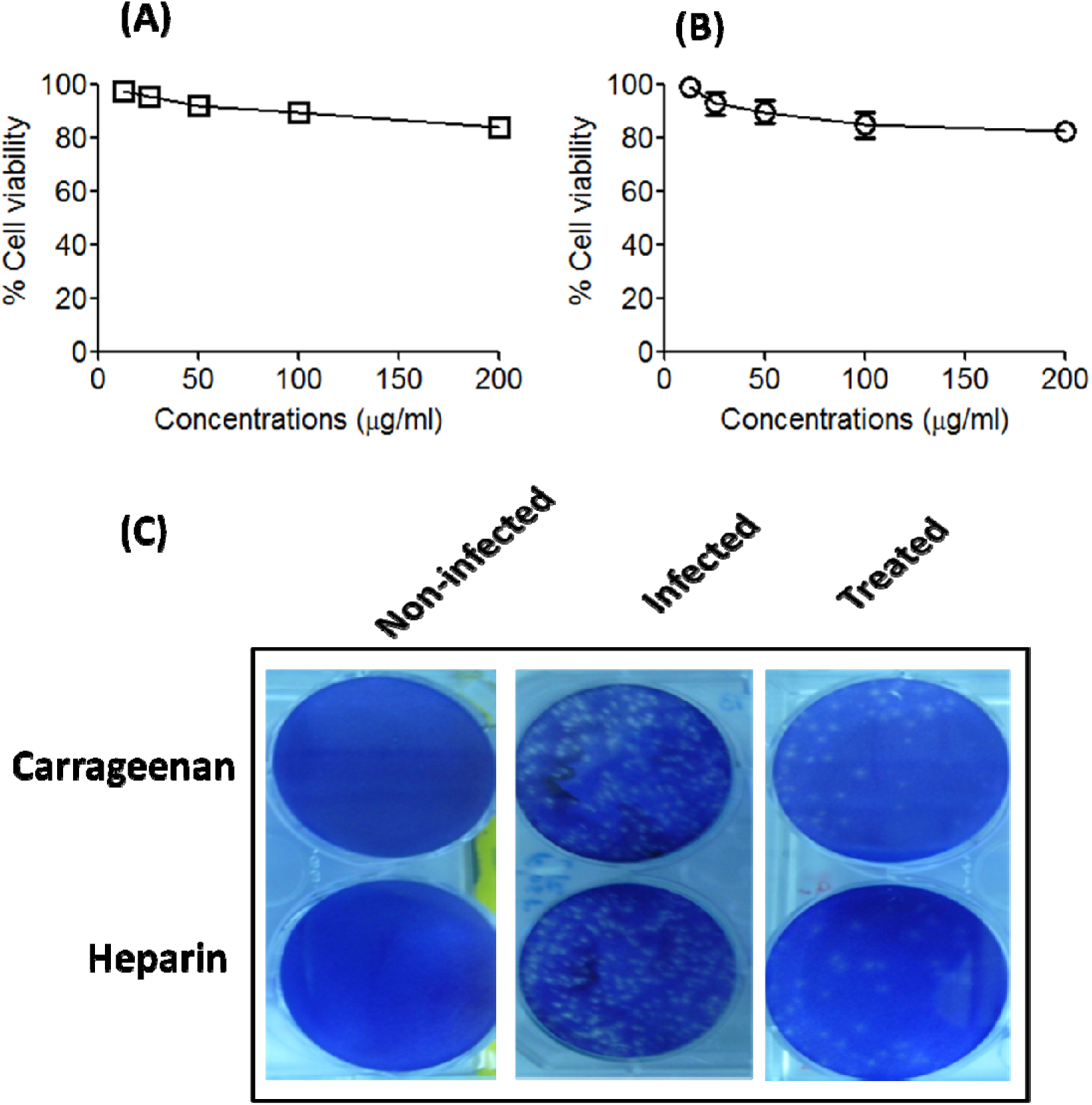
Cell viability of WRL68 cells after treatment with carrageenan and the positive control, heparin and test the antiviral activity by plaque assay. **(A)** The CC_50_ value concentration was higher than 200 µg/ml for carrageenan (**B)** The CC_50_ value concentration was higher than 200 µg/ml for heparin. The MNTD value was approximately at 25 µg/ml for both compounds that showed at least 95% cell viability. **(C)** Determination of viral inhibition by the test compounds using plaque formation assay. The cells were pre-treated separately with 25 µg/ml of each test compounds and infected with JEV at MOI of 1. after 72 hrs, a 10-fold serial dilution of culture supernatant of JEV-infected cells was added to fresh Vero cells grown in 6-well plates (1.2× 10^6^ cells). Viral plaques were stained with crystal violet dye after 5 days of incubation. MNTD=Maximal non-toxic dose.

### Carrageenan inhibited JEV entry into host cells

Liver cell lines (WRL68) showed normal morphology before infection with JEV virus. After JEV-infection, the cells showed various cytopathic effects (CPE). Treatment with carrageenan or heparin showed approximately normal monolayer sheets with lessen CPE (Fig. 2). Then, the antiviral activity of carrageenan against JEV was investigated by plaque formation assay. Our results showed that carrageenan acts by inhibiting viral entry into WRL68 cells, and the inhibitory strength of carrageenan towards JEV was at a similar level as heparin. The results obtained from the direct virus inactivation assay (Fig. 3A) demonstrated that carrageenan had a significant inhibitory effect against JEV (6.7±2.2 p.f.u./ml) compared to untreated cells (19.6±1.8 p.f.u./ml) while no significant difference was observed for heparin as a positive control (7.2±2.6 p.f.u./ml) compared to carrageenan (t-test, p<0.001). Similarly, carrageenan also showed significant inhibitory effects at the viral attachment stage with 4.7±3.1 p.f.u./ml inhibition for JEV infection compared to untreated cells (18.2±2.4 p.f.u./ml) and no significant difference compared to heparin (3.8±2.9 p.f.u./ml) as a positive control (t-test, P<0.01) as presented in Figure 3B. We then incubated the WRL68 cells with JEV at 37°C for two h and subsequently treated the cells with the test compounds to determine their inhibitory effects against viral replication post-infection. The post-infection treatment showed that carrageenan reduced viral titter to 13.4±2.2 p.f.u./ml compared to untreated cells (21.7±2.6 p.f.u./ml). For post-infection assay heparin had showed similar inhibition towards JEV (13.2±3.8 p.f.u./ml) infection (t-test, P<0.05) (Fig. 3C). Moreover, treatment with carrageenan also showed significant protective effects which retained the viability of infected cells to approximately 80% (t-test, p<0.01) compared to untreated cells that showed cell viability of 40% (Fig. 3D)

**Figure 2.**
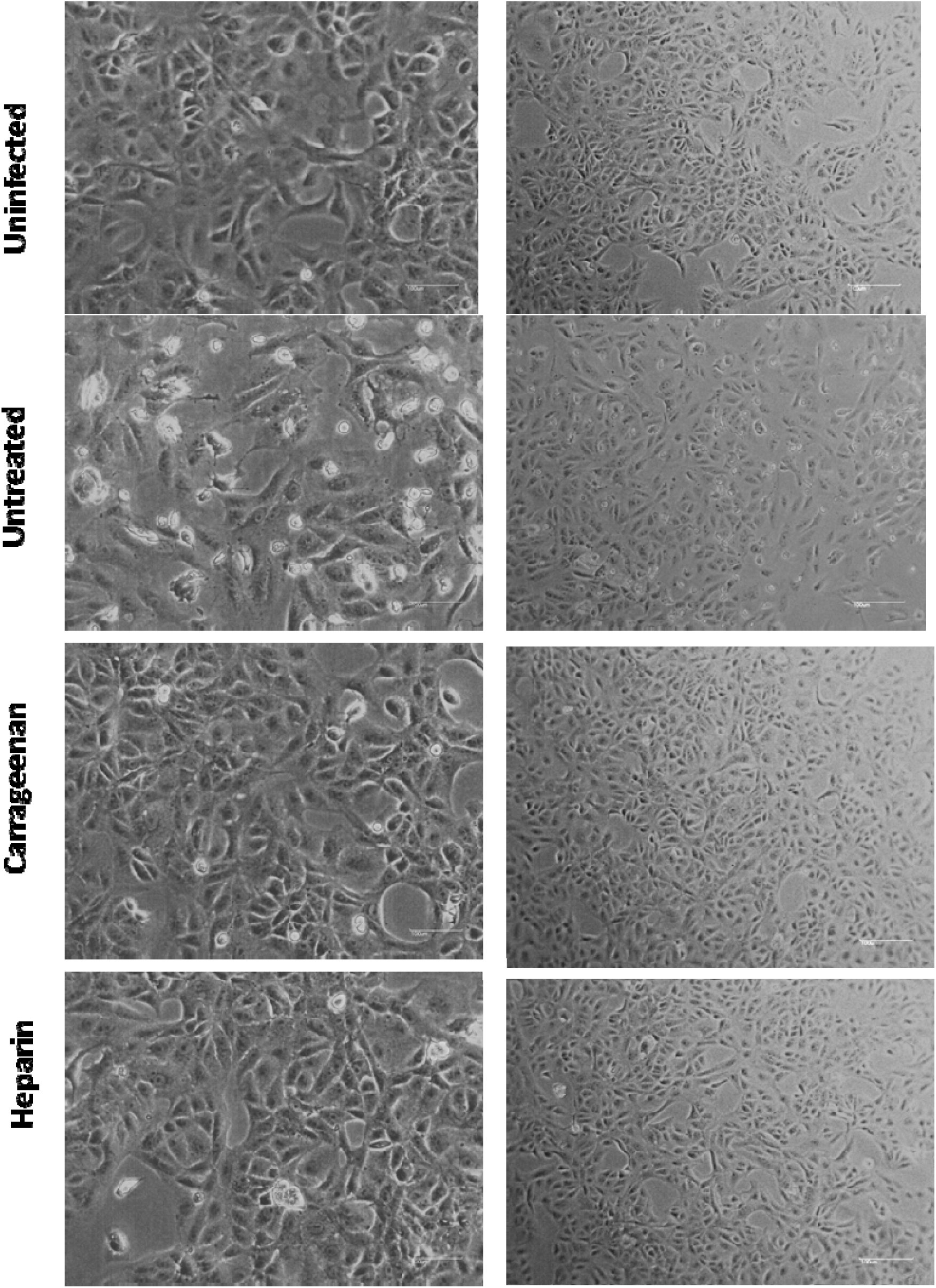
Morphological changes of JEV-infected cells treated with Carrageenan or heparin. Uninfected cells showed normal cell morphology of normal liver cell lines WRL68. The untreated cells morphology post-infection showed various cytopathic effects (CPE). Treatment with carrageenan or heparin showed approximately normal monolayer sheet with lessen CPE. Cell morphology was examined using an inverted microscope at 200× magnification.

**Figure 3:**
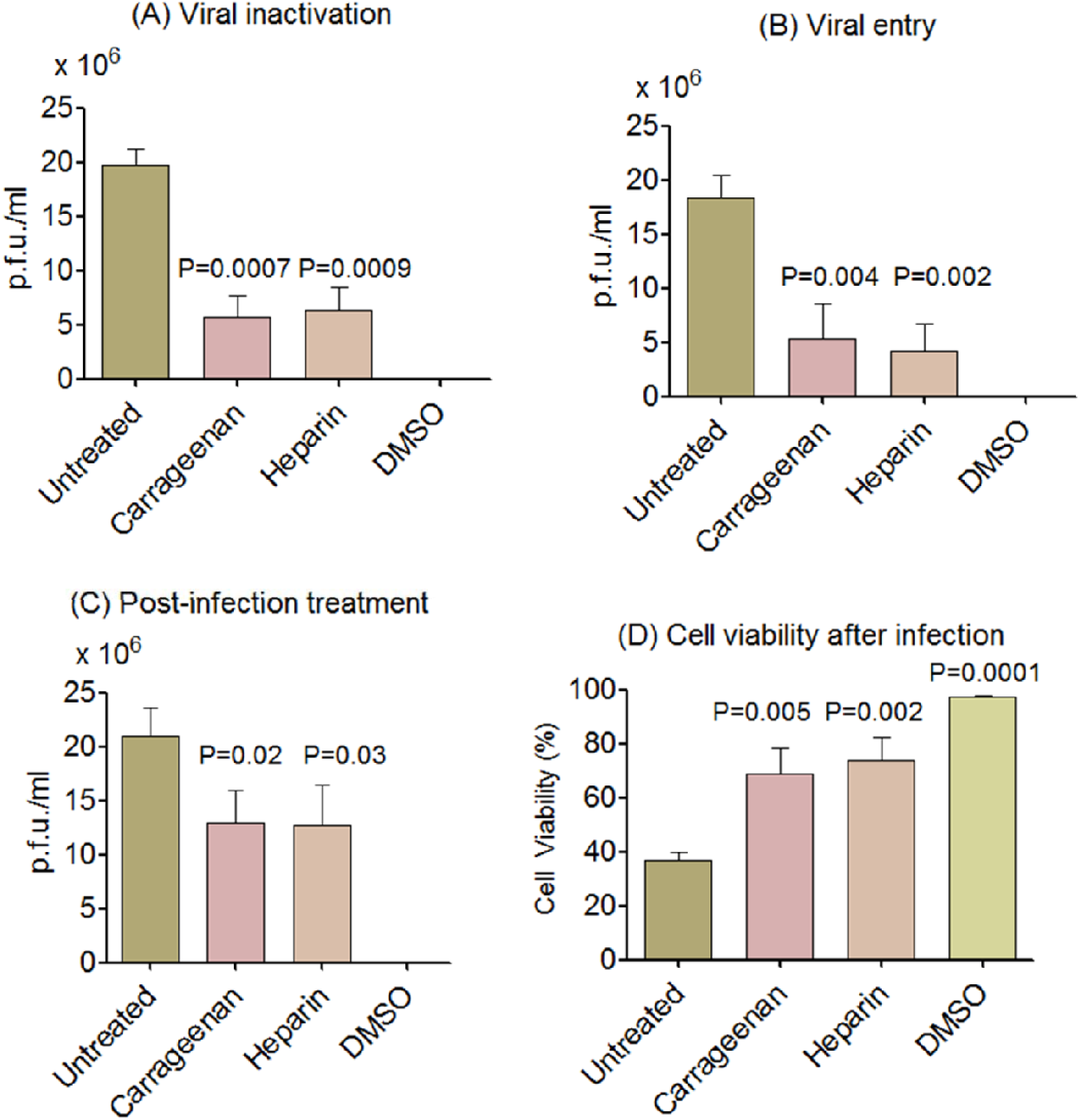
Mode of inhibition of carrageenan against JEV replication. **(A)** Viral inactivation assay showed significant viral inhibition when viral particles were treated with carrageenan compared to untreated cells. Carrageenan showed similar inactivation potential compared to the positive control heparin. **(B)** Viral attachment assay showed that carrageenansignificantly inhibited virus attachment to target cells that was similar to heparin. **(C)** Post-infection treatment showed that the inhibitory effect of carrageenan against JEV was significant but at lower extent compared to the antiviral activity against viral entry (t-test p<0.01). Moreover, treatment with carrageenan also showed significant protective effects which retained the viability of infected cells to more than 75% (t-test, p<0.01).

### Carrageenan inhibited JEV in a dose-dependent manner

Our investigations showed Carrageenan increased inhibition potentials against JEV; we next characterized the antiviral activity of Carrageenan against JEV using an ELISA-like cell-based assay (Fig. 4). Our results demonstrated that Carrageenan inhibited JEV replication in WRL68 cells in a dose-dependent manner. The 50% effective concentration (EC50) for Carrageenan was approximately 15 µg/ml, which is similar to the positive control heparin (Fig. 4). The immunostaining experiment showed that the concentration level of virus particles was considerably decreased after applying increased concentrations from 10 µg/ml to 30 µg/ml of Carrageenan (Fig. 5).

**Figure 4:**
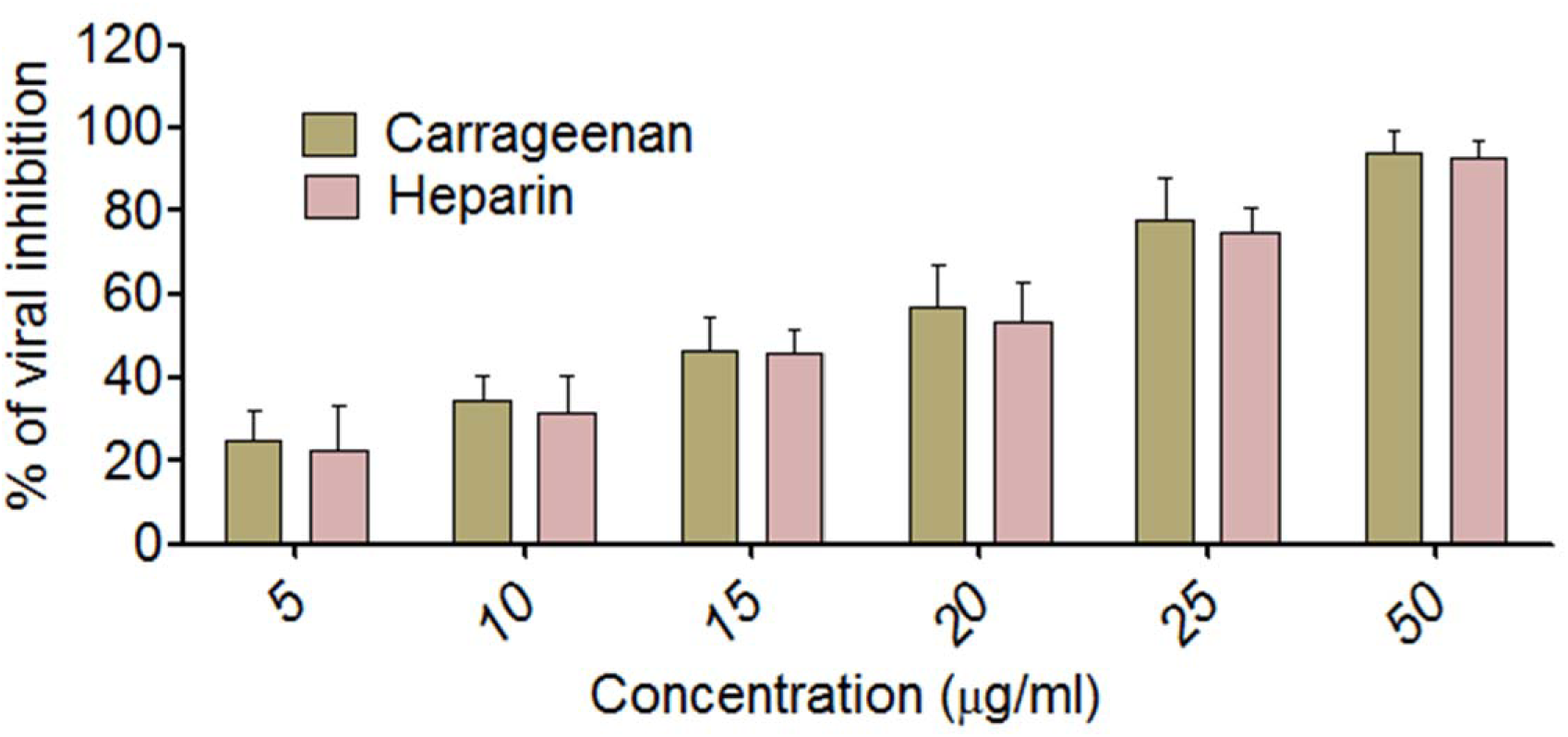
Viral inhibition of JEV in WRL68 cells by carrageenan. Carrageenan inhibited JEV replication in WRL68 cells in a dose-dependent manner. The EC_50_ value for carrageenan was approximately 15 µg/ml which is approximately similar to the positive control heaprin.

**Figure 5.**
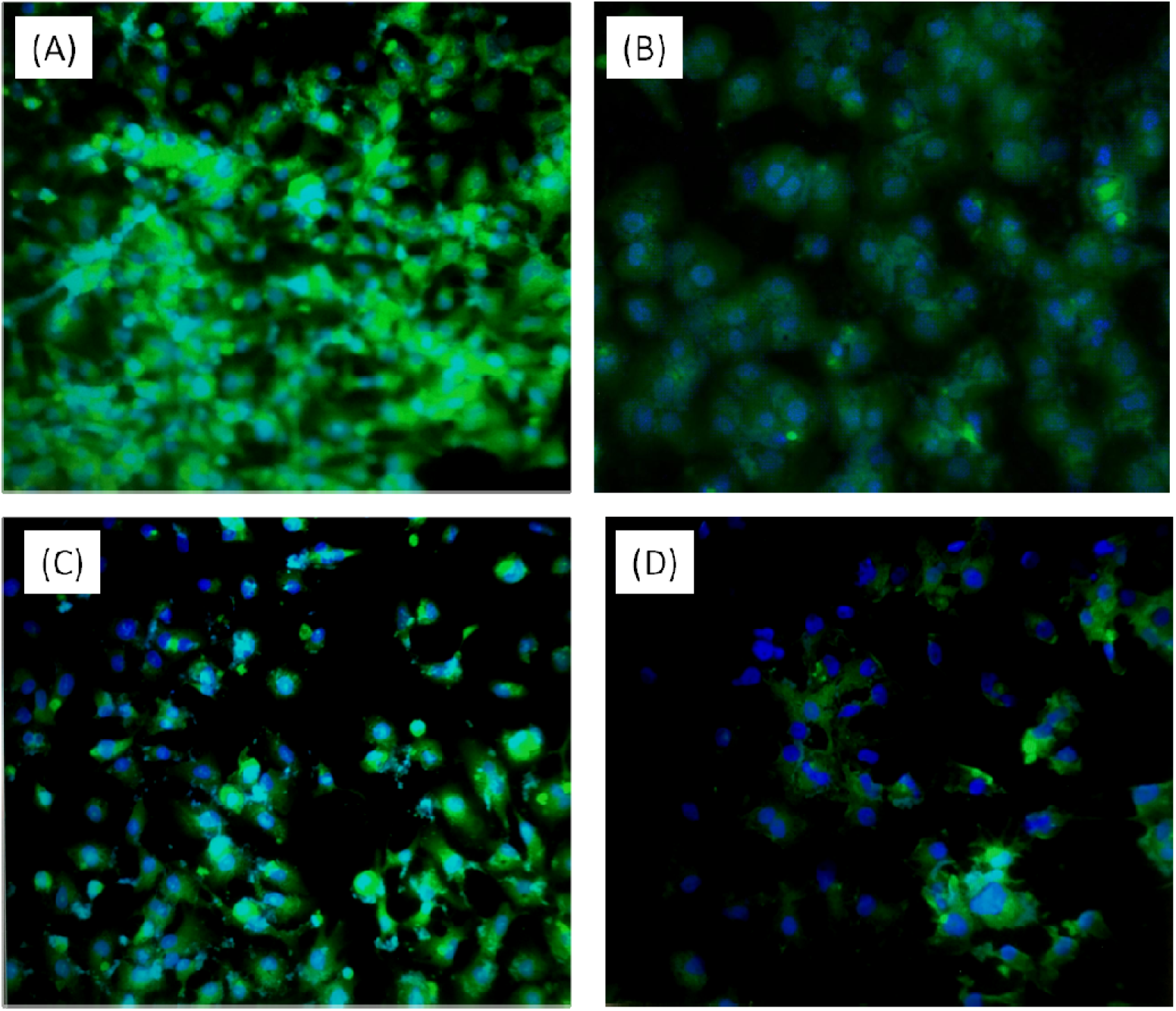
The effect of carrageenan in reducing viral particles. The cells were grown on cover slips in 6-well plates and infected with JEV. The cells were then treated with increasing concentrations of carrageenan (**A**, 0 µg/ml; **B**, 10 µg/ml; **C** 20 µg/ml; **D**, 30 µg/ml) for 72 h. The fixed cells were treated with anti-JEV mouse antibodies and then treated with fluorescein (FITC)-labeled goat anti-mouse IgG. The results showed that the inhibitory effects were dependent on increasing concentrations of carrageenan.

## Discussion

In response to the urgent need for effective anti-JEV drugs, this study was proposed to identify and evaluate new inhibitors for the treatment of JEV infections. Best of our knowledge, the potential of the use of Carrageenan to inhibit JEV has not been previously investigated. Our studies have indeed shown that Carrageenan demonstrated minimal cytotoxicity while exhibiting significant inhibitory effects towards JEV (EC50 = 15 µg/ml) infections in the normal human liver cell (WRL68), suggesting the potential of Carrageenan as an antiviral agent to combat JEV infections. In all experiments, Heparin was used as a positive control.

In this study, three assays were performed to elucidate the antiviral mechanisms of Carrageenan. Our results indicate that Carrageenan inhibits JEV mainly through its direct virucidal effects and interference within the early stages of viral infection, such as viral attachment and cellular entry. We postulated that the direct antiviral activity of Carrageenan might be due to their ability to interact with the virus envelop glycoprotein, particularly the flavivirus E protein, which is responsible for JEV adsorption through the host membrane [18, 19]. Besides, Carrageenan also showed a significant inhibitory effect in the viral attachment assay. In contrast to the direct virucidal activity, the prevention of viral attachment might be attributed to the ability of Carrageenan to compete with both viruses for the cellular receptors (i.e. heparin sulphate) required for cellular entry and subsequent infections [18, 20].

On the other hand, in the post-infection assay, Carrageenan showed lower but significant antiviral activities towards JEV infection. It has been found that heparan sulphate (HS) is involved in the first interaction with E glycoprotein to initiate a flavivirus multiplication cycle in different types of vertebrate cells [21]. HS is a member of highly sulphated glycosaminoglycans (GAGs), which is very abundant on the surface of most mammalian cells and serves as a receptor for many microbial agents, including bacteria, parasites, and viruses [22]. For DENV, the interaction with HS is unusual, owing to its specificity for a highly sulphated form of HS [23]. Therefore, the polysaccharide, curdlan sulphate showed a potential antiviral activity through interfering with the cellular entry of DENV [24].

Our studies also showed that Carrageenan exhibited differing levels of antiviral strength towards JEV, in each of the assays that have been carried out. Although the underlying reasons are still unclear, we suspect it may be due to the variability in the viral glycoproteins and cellular receptors, which are crucial for each virus to establish infection. On the other hand, Heparin has shown similar inhibitory activity in the post-infection assay. Although the potential use of Carrageenan to inhibit JEV, there are still some limitations that have not been addressed in this study. Researches on polysaccharides as antivirals has shown that the antiviral properties of polysaccharides are highly dependent on the structural complexity (i.e. number and position of the branches, branch length and chemical modifications of the polysaccharides) [25, 26]. However, it is difficult for researchers to accurately characterize and purify a particular type of polysaccharide from the extracts of natural sources, which also remains as an inherent limitation in this study. To this end, further research is still needed to elucidate the exact mechanism of inhibition by Carrageenan and its association with the JEV virus replication cycle. Downstream studies with suitable animal models are perhaps required to evaluate the reproducibility of the inhibitory effects of Carrageenan *in vivo*.

In conclusion, our study has successfully demonstrated the inhibition of JEV replication by Carrageenan with minimal cytotoxicity effects. Carrageenan could potentially be a promising broad-spectrum antiviral agent targeting the flavivirus family, and thus warrants further investigation.

